# Mapping morphological malformation to genetic dysfunction in blood vessel organoids with 22q11.2 Deletion Syndrome

**DOI:** 10.1101/2021.11.17.468969

**Authors:** Siyu He, Cong Xu, Yeh-Hsing Lao, Shradha Chauhan, Yang Xiao, Moshe J. Willner, Yinuo Jin, Shannon McElroy, Sneha B. Rao, Joseph A. Gogos, Raju Tomer, Elham Azizi, Bin Xu, Kam W. Leong

## Abstract

DiGeorge Syndrome, or 22q11.2 deletion syndrome (22q11.2 DS), is a genetic disorder caused by microdeletions in chromosome 22, impairing the function of endothelial cells (EC) and/or mural cells and leading to deficits in blood vessel development such as abnormal aortic arch morphology, tortuous retinal vessels, and tetralogy of Fallot. The mechanism by which dysfunctional endothelial cells and pericytes contribute to the vasculopathy, however, remains unknown. In this study, we used human blood vessel organoids (VOs) generated from iPSC of 22q11.2 DS patients to model the vascular malformations and genetic dysfunctions. We combined high-resolution lightsheet imaging and single-cell transcriptome analysis to link the genetic profile and vascular phenotype at the single-cell level. We developed a comprehensive analytical methodology by integrating deep learning-mediated blood vessel segmentation, network graph construction, and tessellation analysis for automated morphology characterization. We report that 22q11.2DS VOs demonstrate a smaller size with increased angiogenesis/sprouting, suggesting a less stable vascular network. Overall, clinical presentations of smaller vascular diameter, less connected vasculature, and increased branch points were recapitulated in 22q11.2DS VOs. Single-cell transcriptome profiling showed heterogeneity in both 22q11.2DS and control VOs, but the former demonstrated alterations in endothelial characteristics that are organ-specific and suggest a perturbation in the vascular developmental process. Intercellular communication analysis indicated that the vascular dysfunctions in 22q11.2 deletion were due to a lower cell-cell contact and upregulated extracellular matrix organization involving collagen and fibronectin. Voronoi diagram-based tessellation analysis also indicated that the colocalization of endothelial tubes and mural cells was different between control and 22q11.2 VOs, indicating that alterations in EC and mural interactions might contribute to the deficits in vascular network formation. This study illustrates the utility of VO in revealing the pathogenesis of 22q11.2DS vasculopathy.

## Main

22q11.2 deletion syndrome (22q11.2 DS) results from a microdeletion of up to 50 genes on the long arm of chromosome 22. Patients experience variable developmental dysfunction, such as abnormal aortic arch morphology, congenital heart disease, and neuropsychiatric disorders^1-4^. Previous studies have reported that the vascular malformations in 22q11.2 DS are mainly due to the haploinsufficiency of *T-box Transcription Factor 1* (*TBX1), DiGeorge Critical Region 8 (DGCR8)*^*4*^, ^*5*^, affecting VEGF-associated signaling pathways, such as *DLL4/Notch1-VEGFR3*^*6*^ and *AKT/*GSK3β/VEGF signaling pathways^7^, ^8^. Downstream pathological phenotypes of vascular malformation and dysfunction include hyperplasia, enhanced angiogenic spouting, network disorganization, defect vessel perfusion, and hyperoxia^6^, ^9-11^. Although malformations have been identified in individuals with 22q11.2DS, the lack of robust quantification hinders the diagnostic standardization and deeper understanding of its derived vascular malformation.

Recent advances in stem cell engineering enable in vitro generation of three-dimensional (3D) self-organized tissues, known as organoids, with either embryonic stem cells (ESCs) or human induced pluripotent stem cells (hiPSCs). These models partially recapitulate human physiology and they have emerged as transformative toolkits for disease modeling and drug screening. The recent development of hiPSC-derived blood vessel organoids (VOs) is such an example. It resembles human blood vessels with respect to cell composition (e.g., presence of both endothelial cells and pericytes) and 3D vascular architecture (e.g., tubular-like structures enveloped by the basement membrane)^12^. Transplantation of these VOs in diabetic mice produces microvascular changes observed in diabetic patients such as vessel leakiness and thickening of the vascular basement membrane^12^. Recently, the VOs have also proved a valuable model in understanding if human recombinant soluble ACE2 (hrsACE2) can inhibit SARS-CoV-2 infections^13^.

Although VOs have been applied in a number of disease models, most studies focus on qualitative analyses. Quantitative analysis of 3D VO morphology remains challenging due to the limited capacity of manual analysis, which is time-consuming, laborious, subjective, and biased toward a selected region of interest (ROI). Recent advances in deep learning technology enable automated accurate 3D segmentation of blood network structure, which provides reliable objects for the downstream analysis to reveal the pathological phenotype changes during disease progression^14^. Here, we generated 84 VOs in total from three pairs of 22q11.2 DS patients and normal controls iPSC lines (Fig. 1a, Fig. S1a). we applied deep learning techniques to characterize the vasculature malformation in 22q11.2 DS including the changes in vascular diameter and network connectivity. We further established an integrated analysis pipeline to map the morphological changes to single-cell RNA expression alterations to study the molecular basis of dysfunction in the 22q11.2 DS VOs (Fig.1b). Our single-cell transcriptomic analyses revealed abnormalities in both vascular and vascular-associated cell types in cell-cell contact signaling and basement malformation in the 22q11.2DS vasculature. Finally, a Voronoi diagram-based tessellation analysis demonstrated a weaker colocalization of endothelial tubes and mural cells suggesting a consistent trend in decreased cell-cell contact. Taken together, this study establishes a platform that integrates the merits of tissue engineering, deep learning, graphic analysis, single-cell sequencing to study some key features of vascular cells and developing vascular networks. With this platform we revealed some important abnormalities in the pathogenesis of 22q11.2 deletion vasculopathy; this platform should also be applicable to other vascular disease modeling studies.

**Figure 1.**
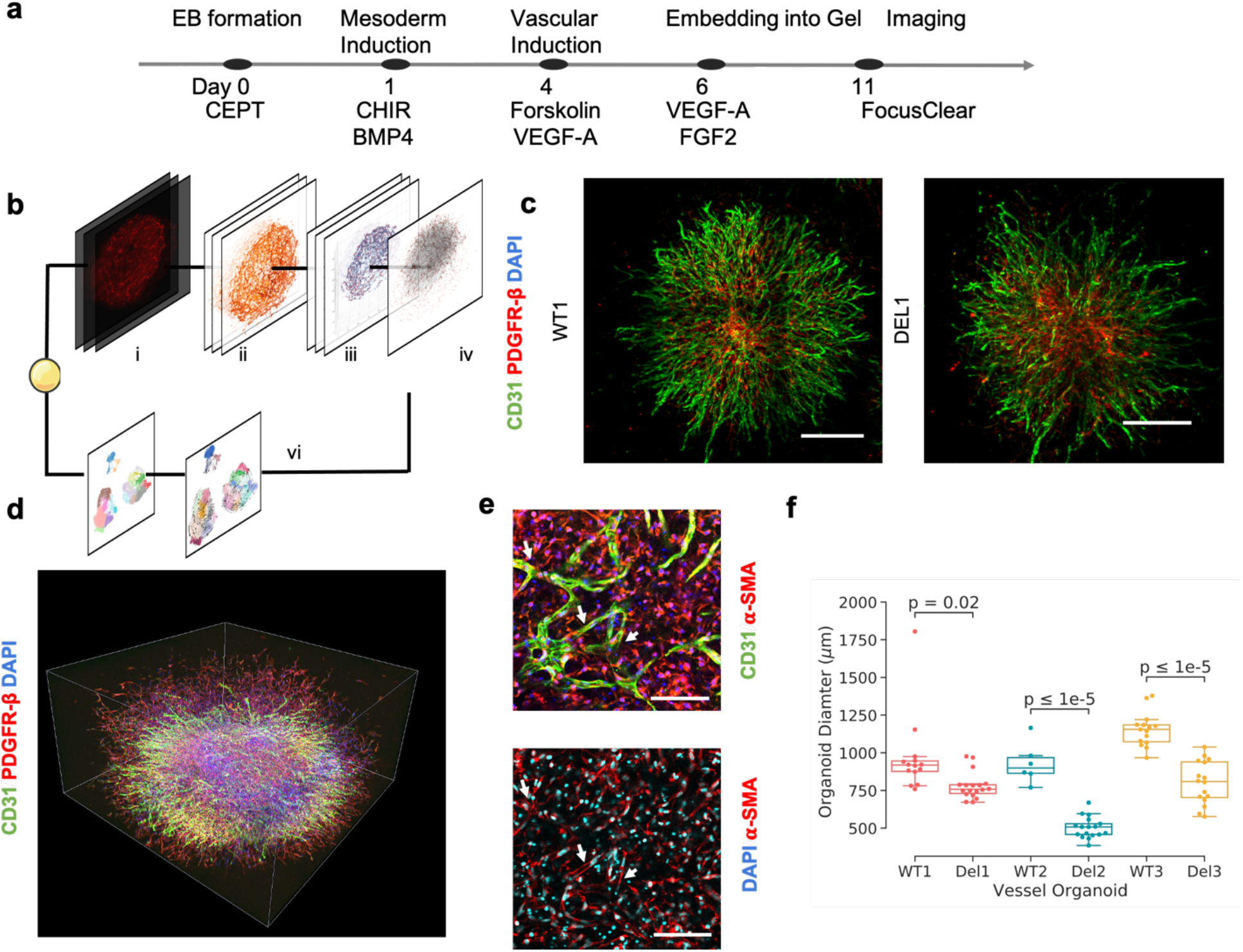
Experimental and analytical schematics of blood vessel organoids. a) Protocol of blood vessel organoids. b) Schematic of the workflow combining the quantitative analysis of vasculature phenotype and single cell transcriptome. i: COLM imaging; ii: 3D segmentation by deep learning; iii: Morphology features; iv: topology features; vi: single cell analysis including trajectory inference and intercellular communications. c) Exemplary Z projection of the blood vessel organoid derived from one paired iPSC line (WT1 and Del1, Scale bar: 200 um). d) 3D rendering of one exemplary blood vessel organoid, with the staining of CD31, PDGFR-β, and DAPI. e) Zoom in of EC and mural cells in VO (Scale bar: 300um). f) Comparison of organoids size in 22q11.2DS vessel organoids and control (Mann-Whitney-Wilcoxon test, *p* = 2.142×10^−2^, 6.352×10^−9^, 7.167×10^−8^ and fold change = -0.19, -0.45, -0.30, respectively; *n* = 86 in total).

## Results

### Generation and characterization of blood vessel organoids

We first followed the previously established protocols to generate VOs^12^, ^15^ (Fig. 1a). At D11, VOs were fixed and immunostained for 3D-mount lightsheet imaging (Fig. S1b) or subjected to single-cell RNA sequencing for further integrated analysis (Fig.1b). Through staining of the endothelial cell (EC) tubes (CD31^+^) and mural cells (PDGFR-β_+_, α-SMA^+^), the VOs showed an architecture similar to what was reported in the literature^12^. The Z-projection of generated 3D VO image stacks between the 22q11.2 DS and control groups both showed the recapitulation of vascular network constructed by endothelial tubes and mural cells (Fig. 1c, supplementary video). The 3D imaging of VOs indicated the complexity of the network structures and 3D alignment of the EC and mural cells (Fig.1de, and Supplementary Video). For example, as shown in Fig. 1e, the EC tubes aligned with the mural cells. These results indicated that our VOs harbored the capability of mimicking vascular network morphology in both healthy and 22q11.2DS patient blood vessels.

### Morphological alterations in 22q11.2 DS blood vessel organoids

To characterize the morphological variation in 22q11.2 VO, we first measured organoid size. The VO volume derived from three 22q11.2 DS patient iPSC lines was smaller than their control counterparts (Fig. 1f). This result is consistent with the clinical findings that 22q11.2 DS patients have reduced total brain volume (correlated with the cerebrovascular lesions^16-18^), and deficit in vascular development^19^, ^20^. The heatmap of DAPI staining in the midplane of the 22q11.2 Vos showed alterations of cell density distribution (Fig. S1cd). These findings suggest there are cellular proliferation and/or migration differences between 22q11DS and control VOs.

To quantitatively determine the tubular structure in the VOs, we first modified a fully convolutional neural network (FCN) deep learning architecture, namely DeepVesselNet^14^, ^21^, which has been trained to study vascular networks, to detect and segment the EC tubes in VOs. Convolutional neural network (CNN) has been widely applied in the field of image recognition due to the ability to identify both concrete, such as shapes and edges, and abstract characteristics of the images. The architecture consists of four CNN layers, one fully connected layer, and a sigmoid activation^21^. The output of architecture includes predicted probability in the 3D microvascular space. The extracted endothelial network was visualized with its vessel radius distribution, and the tubular shapes and sprouting of the microvasculature were well-maintained in the images throughout the whole organoid (Fig. S1d). To reveal the density variation of endothelial tubes among the 3D space and cells, we measured the density distribution of vascular microtubes based on the 3D segmentation, followed by a Gaussian kernel density estimation^22^ for the entire VO. Fig. 2a presents the representative density distribution (normalized to 1.0) of 22q11.2 DS and control VOs (termed Del and WT1, respectively). In two patient lines of 22q11.2 DS VOs, high-density vascular microtubes (density distribution > 0.95) occupied a larger 3D vascular volume than those in the control VOs (Fig. S1e). One patient/control pair did not show significant difference (WT2 and Del2), which might be due to the variability of the cell line; nevertheless, the mean vascular density of all three patient-derived VOs is a significantly higher value compared to that of their control counterparts (Fig. 2b). To determine whether vascular density distribution was correlated with cell distribution, we performed nuclei segmentation by implementing CellPose^23^ on the same organoids. The sum of nuclei number over Z-dimensions (Fig. 2c) showed an even distribution of nuclei across samples, indicating that the nuclear density and vascular density of the same VO were not linearly correlated. With regard to the number of nuclei, in all of our organoids, 22q11. 2DS VOs had a lower total nuclei number (Fig. 2b). These results suggest that the higher density of vascular network in 22q11.2 is not due to a higher number of cells. Instead, it is associated with specific features of dense vascular tubes. To determine which features contribute to the higher density of vascular network in 22q11.2 DS VOs, we carried out a global statistical analysis on vascular features for each sample. All the results were calibrated from pixel volume to physiological space. The mean and maximum vascular radii of 22q11.2 DS VOs (r_mean_ = 8.03, 6.83, 7.56 µm, r_max_ = 23.17, 21.18, 22.12 µm for Del1, Del2, Del3, respectively) were significantly lower than that in the control VOs (r_mean_ = 8.06, 8.02, 8.15 µm, r_max_ = 31.29, 33.17, 31.87 µm for WT1, WT2, WT3, respectively; *p* = 7.87×10^−1^, 1.10×10^−2^, 1.63×10^−3^ with Mann-Whitney-Wilcoxon test, fc = -0.005, -0.148, -0.073; *n* = 84 in total). The magnitude of the vascular radius in the generated VOs was close to that of human blood vessel^24^, demonstrating that the blood vessel structures in our VO were representative. In a comparison of the mean and maximum lengths of the EC tubes, the 22q11.2 DS VOs did not show a significant difference as compared with the controls (Fig. S1f). The density and mean radius are also dependent on the distance from the VO center (Fig. S2ab). Furthermore, we analyzed the skeletonization in each sample (Fig. S1g). Three types of bifurcation point types in the organoid were determined by the number of branches linked to them (junction points, path points, and endpoints, Fig. 2d). Enrichment of endpoints was found in the 22q11.2 DS VOs, indicating the enriched tubular sprouting compared with the controls (Fig. 2b), while junction points and path points didn’t show significant differences (Fig. S1f). The alterations in the morphological phenotypes indicated the difficulty of forming large blood vessel organoids in 22q11.2DS VO, though the vascular density or the ability of angiogenesis/sprouting increased.

**Figure 2.**
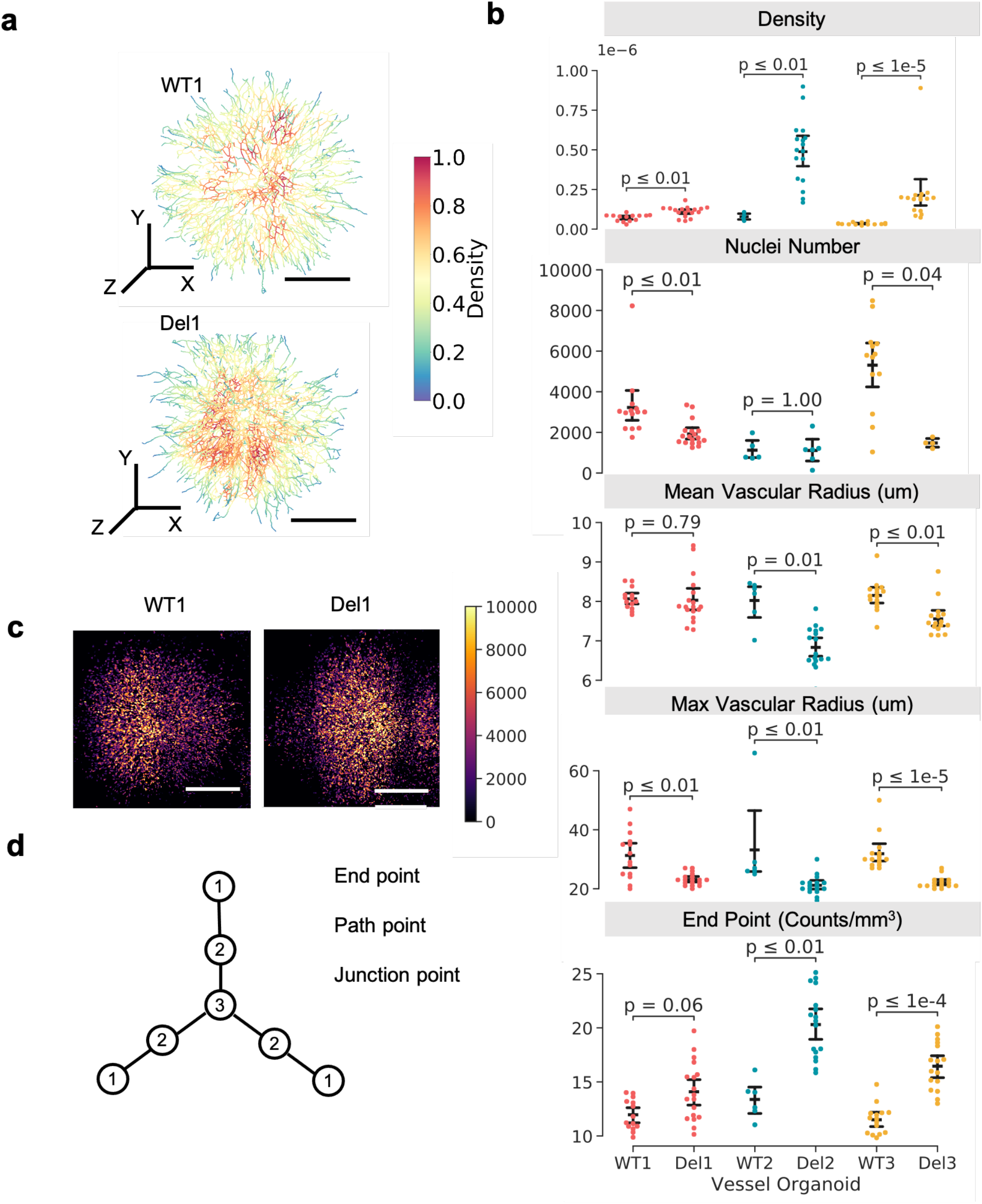
Morphological variations of 22q11.2DS vessel organoids segmentation via deep learning. a) 3D rendering of control and 22q11.2DS VO, colored by the density value of vascular tubes. The density in each VO is normalized to [0,1]. Scale Bar: 200 um. b) Statistical comparison of vascular density, radius, nuclei number, and endpoint of blood vessel structure in three pairs of VOs (Mann-Whitney-Wilcoxon test, Density: *p* = 3.4×10^−3^, 8.1×10^−3^, 6.9×10^−6^, fc = 0.54, 5.22, 5.19; Nuclei number: *p* = 1.15×10^−3^, 1, 4.48×10^−2^, fc = -0.40, -0.01, -0.72; Mean vascular radius: r_mean_ = 8.03, 6.83, 7.56 µm, for Del1. Del2, Del3; r_mean_ = 8.03, 6.83, 7.56 µm for WT1, WT2, WT3; *p* = 7.87×10^−1^, 1.10×10^−2^, 1.63×10^−3^, fc = -0.005, -0.148, -0.073; Max vascular radius: r_max_ = 23.17, 21.18, 22.12 µm, for Del1. Del2, Del3; r_max_ = 23.17, 21.18, 22.12 µm, for WT1, WT2, WT3; *p* = 7.87×10^−1^, 1.10×10^−2^, 1.63×10^−3^, fc = -0.005, -0.148, -0.073; End point: *p* = 6.465×10^−2^, 1.587×10^−3^, 1.803×10^−5^; fc = 0.164, 0.518, 0.431). c) Z projection of nuclei staining of the control and 22q11.2DS VO. d) Diagram of network and classification of branch points (end point, path point, junction point).

### Topological alteration identified among 22q11.2 DS vascular graphs

To understand the structural properties of the blood vascular networks in VO, we investigated the geometric and topological features, such as tortuosity, clustering of vascular nodes or inter-node connectedness, by constructing a network graph for each VO image. This topological vessel model is known to correlate with biological processes such as blood vessel flow and cell-cell communications^25^. Endpoint and junction points in Fig.2d were considered as the nodes, and vascular branches were considered as the edges that connect the nodes in the network graphs. Two representative network graphs obtained from the 22q11.2 DS and control VOs are shown in Fig. 3a; red dots and grey lines denote the nodes and the edges, respectively. We first investigated the connectivity, which is defined as the ratio of the joint points over the endpoints. The 22q11.2 DS VOs had a lower connectivity as compared with the controls (Fig. 3b). The finding suggests that the sprouting ability in the 22q11.2 DS VOs is significantly enhanced. We performed the tortuosity measurements based on the absolute distance of vascular branches dividing the Euclidean distance between nodes. As shown in Fig. S2c, there was no significant difference in the tortuosity and clustering coefficient between all three 22q11.2 DS samples and the controls. Subsequently, we investigated the degree distribution *P*(*k*) of the vascular nodes in VOs (Fig. 3c). The relative frequency of degree in the diseased case (blue line) showed a higher value than the controls (red line) (*k* <7), but lower values over the controls when *k* >7. Previous studies reported that the high-degree nodes would be associated with angiogenic hotspots, or tumor periphery^25^. The distribution has been previously discussed to follow the power law, *P*(*k*)∼*k*^−γ^, where γ is the degree exponent. As presented in Fig. 3d, all the degree exponent values of the 22q11.2 DS and control VOs were within the previously reported range of 2 < γ < 3, classified as “scale-free” networks^25^ that are resilient against perturbations and random failures. Despite that, the 22q11.2 DS VOs behaved with a higher degree exponent (*p* = 1.538×10^−2^, 3.040×10^−2^, 1.682×10^−5^). A higher exponent is known to lead to a loss of global connectivity against perturbations. The results of this analysis suggested that a lack of robustness of the vascular network in the 22q11.2DS VOs, rendering them more vulnerable to lose their global connectivity from a potential attack in the context of graph theory. In the context of physiology, vascular network graphs correlate with alteration of cellular communications, blood flow features, global sizes, and atrioventricular septal defect in 22q11.2DS patients. Speculatively, it provides a new concept derived from network theory to explore the complexity of physiological structure and functions^26-30^.

**Figure 3.**
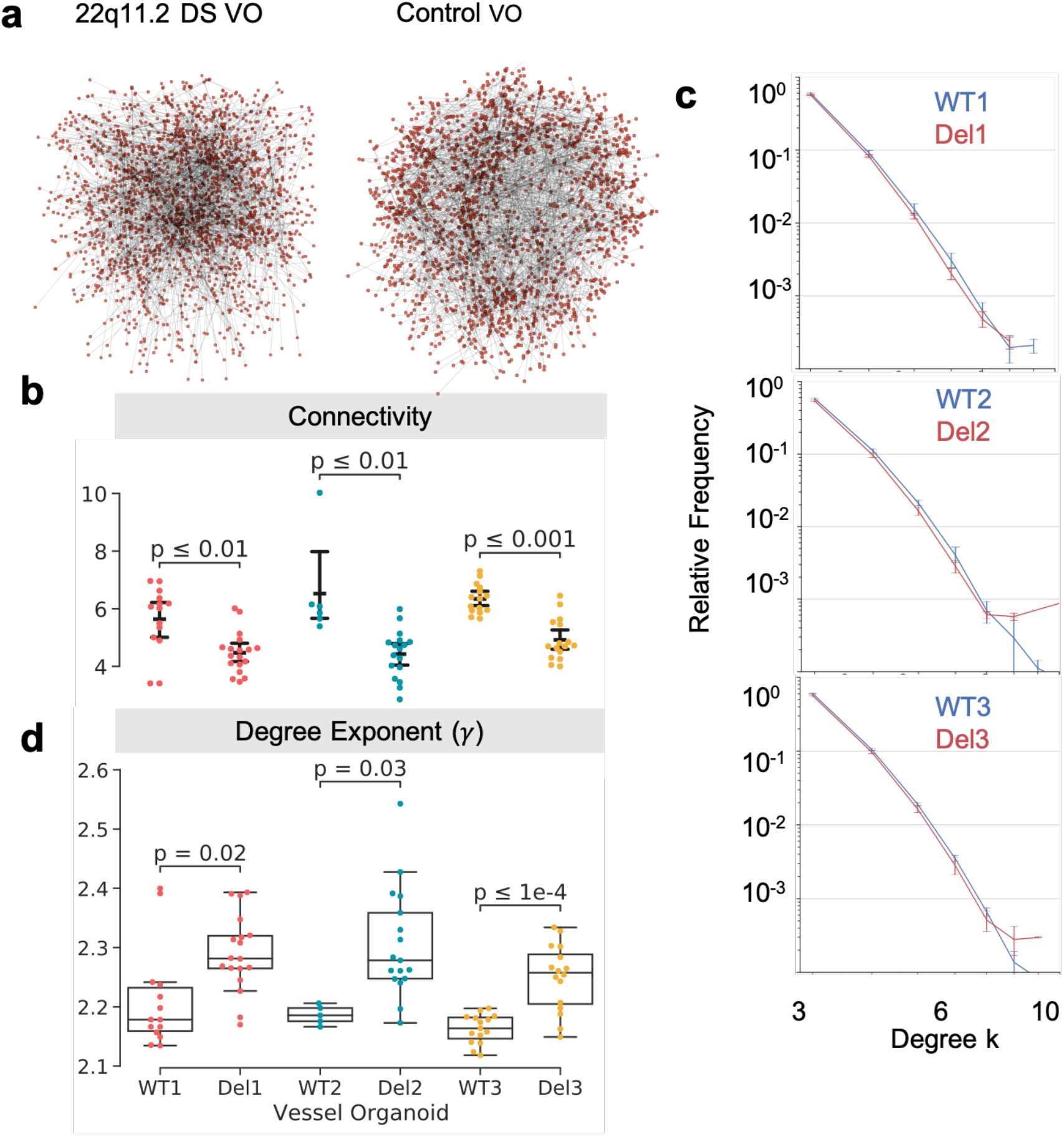
Morphological variations of 22q11.2DS vessel organoids segmentation via deep learning. a) Network graph of 22q11.2 DS VO and control. Red dots denoted graph nodes those derived from the branch junctions, and grey edges denoted connectedness of those nodes respectively. b) Connectivity of 22q11.2 DS and control VO. c) Relative frequency of degree distribution in the vessel organoids graph. d) Degree exponents in the organoids graph (*p* = 8.592×10^−3^, 4.333×10^−3^, 1.526×10^−4^; fc=-0.21, -0.32, -0.22; *n* = 84 in total).

### Colocalization study of endothelial cells and mural cells in 22q11.2 DS VO

From the fluorescent images, we saw colocalization of ECs and mural cells in our VO samples (Fig1e, Fig 4a). To understand the potential physical interaction between these two cell types at the pixel level, we implemented a colocalization analysis based on the Voronoi algorithm^31^ and utilized the two colocalization metrics Manders and Spearman’s coefficients, to evaluate the interactions of EC and mural cells in our VOs. After segmenting VOs with CD31+ and PDGFRb+ at the middle plane (Fig. S3a), we were able to partition the segmentations into polytopes using the Voronoi diagram (Fig. 4b, Fig. S3b). Those Voronoi polytopes were then classified as high-density or background based on the normalized 1^st^ rank density 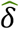 (Fig. S3de)^31^. The localization of one channel could be bound with the other channel based on overlapping polytopes and could be represented with the local pair-normalized localization densities 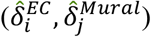 and 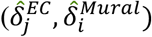 .Manders and Spearman’s coefficients for ECs and mural cells (*M*^*EC*^ *M*^*Mural*^) and (*S*^*EC*^ *S*^*Mural*^) were calculated based on the pair-normalized localization densities. The logarithmic transformed pair-normalized localizations in different conditioned VOs were correlated with the labeled Manders and Spearman’s coefficients (Fig. 4c). To explore the significance of those differences, we performed this Voronoi analysis on all the Z slides of our VOs. As shown in Figs. 4d and 4e, we compared Manders and Spearman Rank coefficients of EC colocalization with mural cells as well as these of mural cell colocalization with EC in the VOs. The 22q11.2 DS VOs had significant decreases in Manders and Spearman Rank coefficients in both EC colocalization and mural cell colocalization. These findings indicated an abnormality of EC and mural co-localization in the 22q11.2 DS samples, suggesting a misalignment of their localization, and/or decreased cell-cell contacts between EC and perivascular cells.

**Figure 4.**
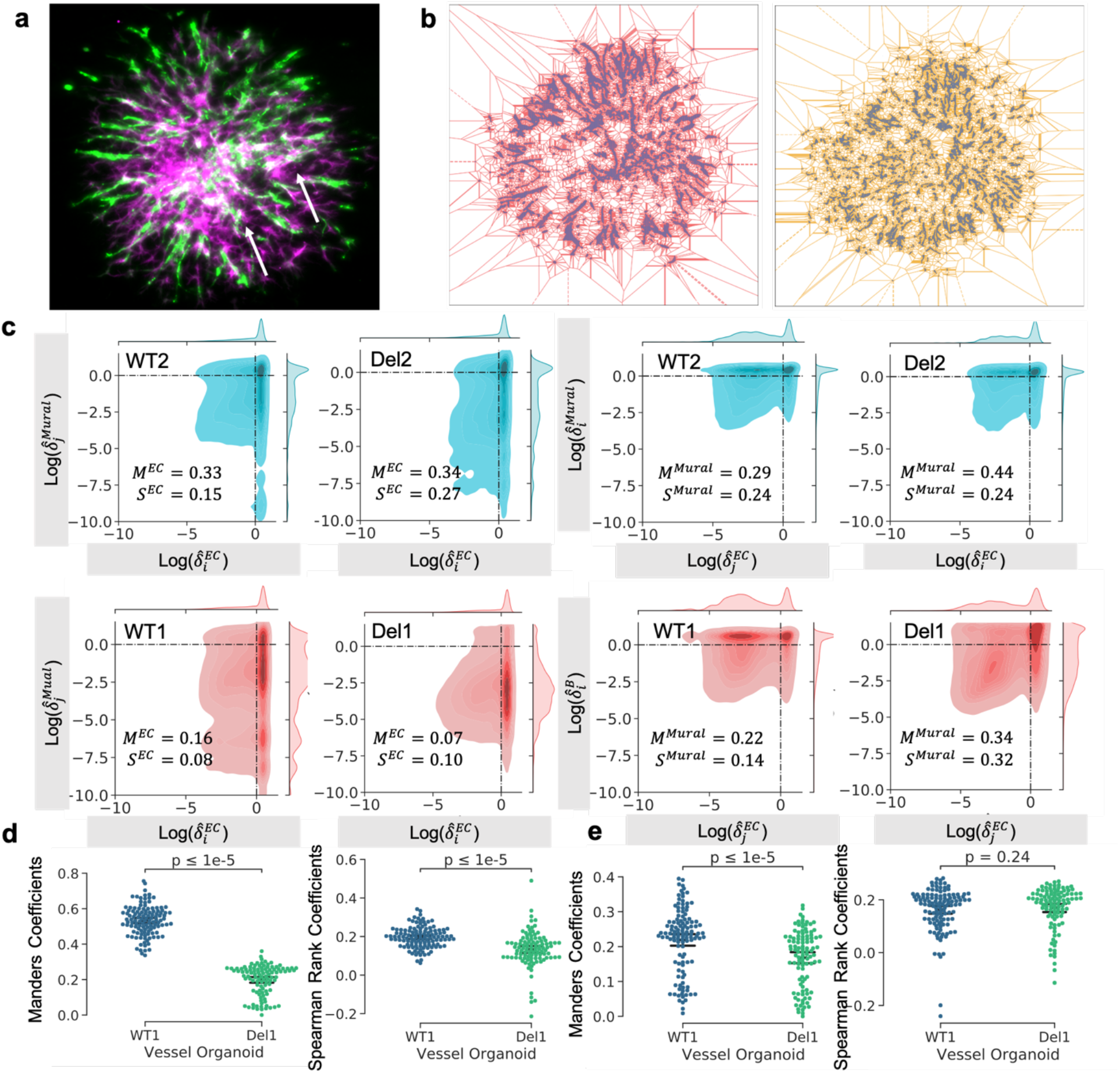
Colocalization analysis of EC and mural cells. a) An example with colocalization of EC and mural cells, where the white arrows referred. b) Voronoi diagram of CD31+ and PDGFRb+ channels. c) Scatterplot of densities pairs of endothelial and mural cells. d) Manders coefficients and Spearman rank coefficients of EC colocalization corresponding to mural cells in VO. e) Manders coefficients and Spearman rank coefficients of mural cell colocalization corresponding to ECs in VO.

### Single-cell RNA profiling of blood vessel organoids

We performed single-cell RNA sequencing on Del3 and WT3 VOs on Day 11 to construct a transcriptome atlas of all cell types within human VOs. We obtained 13,797 cells in total (5,580 cells and 8,217 cells from several WT3 VOs and several Del3 VOs respectively). Principal component analysis was performed after normalization, and the cells from control and 22q11.2DS VO were embedded together in a 2D UMAP space^32^. We identified 25 cell clusters using PhenoGraph (Fig 5a)^33^. Six of those clusters were annotated as ECs with two criteria: First, they have a robust differential expression of EC markers^34^, ^35^ (PECAM1, CDH5, CLDN5, ERG; Fig. S4a); and second, they lack non-EC markers (ACTA2, TNNT2). Six clusters were identified mural cells based on reported marker genes, namely ACTA2, TAGLN, PDGFRβ_34_, and four clusters were identified as fibroblasts based on the differential expression of COL1A1 (Fig. S4a)^34^. Other clusters were annotated as progenitor or proliferating cells based on proliferation marker (Ki-67, PCNA)^36-38^. By applying RNA velocity^39^ to single-cell sequencing data^40^ (Fig.5b), we predicted the trajectory of cell state, from the progenitor cells to endothelial cells or other vascular-associated cells. Trajectories also indicated the differential and developmental process of vascular cells, such as from Mural3/4 to Mural1, Mural5 to Mural2, FB4 to FB1 (Fig. 5b). PAGA^41^ diagrams showed a more general tendency for the vasculature development (Fig. S4b). The pseudo-time map (Fig. 5c) indicated the differentiation from iPSC to mature endothelial cells in both WT3 and Del3 VOs. Based on the gene enrichment and reported gene markers of the cell subtypes, we were able to distinguish arterial (BMX, VEGFC, ARL15, SEMA3G), venous (NR2F2, TFRC, SLC38A5, TSHZ2, ADGRG6, VCAM1), and capillary ECs (MFSD2A, SLC7A5, TFRC, SLC16A1)^35^, ^42^ (Fig. S4cd). Leveraging on the well-characterized EC markers in specific organs, we further determined the organ identity of ECs such as the brain (MFSD2A, SLCO1C1, SLC2A1), kidney (IGFBP5), lung (SCN3B, SCN7A, ACE), and liver (IGFBP5, STAB2, FCGR2B) (Fig. 5d)^34^. With this analysis, we noticed the percentage of the potential kidney-specific EC is dominant in our VOs (59.5% kidney vs. 24% brain or 5.4% lung; Fig. 5e), agreeing with the kidney specificity of the original VO differentiation protocol^12^. However, Bulk RNA sequencing results showed the downregulation of kidney development in the signaling pathway analysis in Del1 VOs supported this finding that ECs are related to the liver-specific cells (Fig S7b). In addition, we specified the mural subtypes by arterial smooth muscle cells markers (ACTA2, MYH11), venous smooth muscle cells markers (MRC1, CD74), and pericytes markers (GRM8, PDGFRB)^42^ (Fig. S4e, Fig. 5f). The heatmap of top differential genes confirmed that each cell type clustered separately (Fig. 5g). Thus, single-cell RNA profiling confirmed the existence of EC and vascular-associated cells, including pericytes, smooth muscle cells, and fibroblast in our VOs, which also indicated the heterogeneity of VOs.

**Figure 5.**
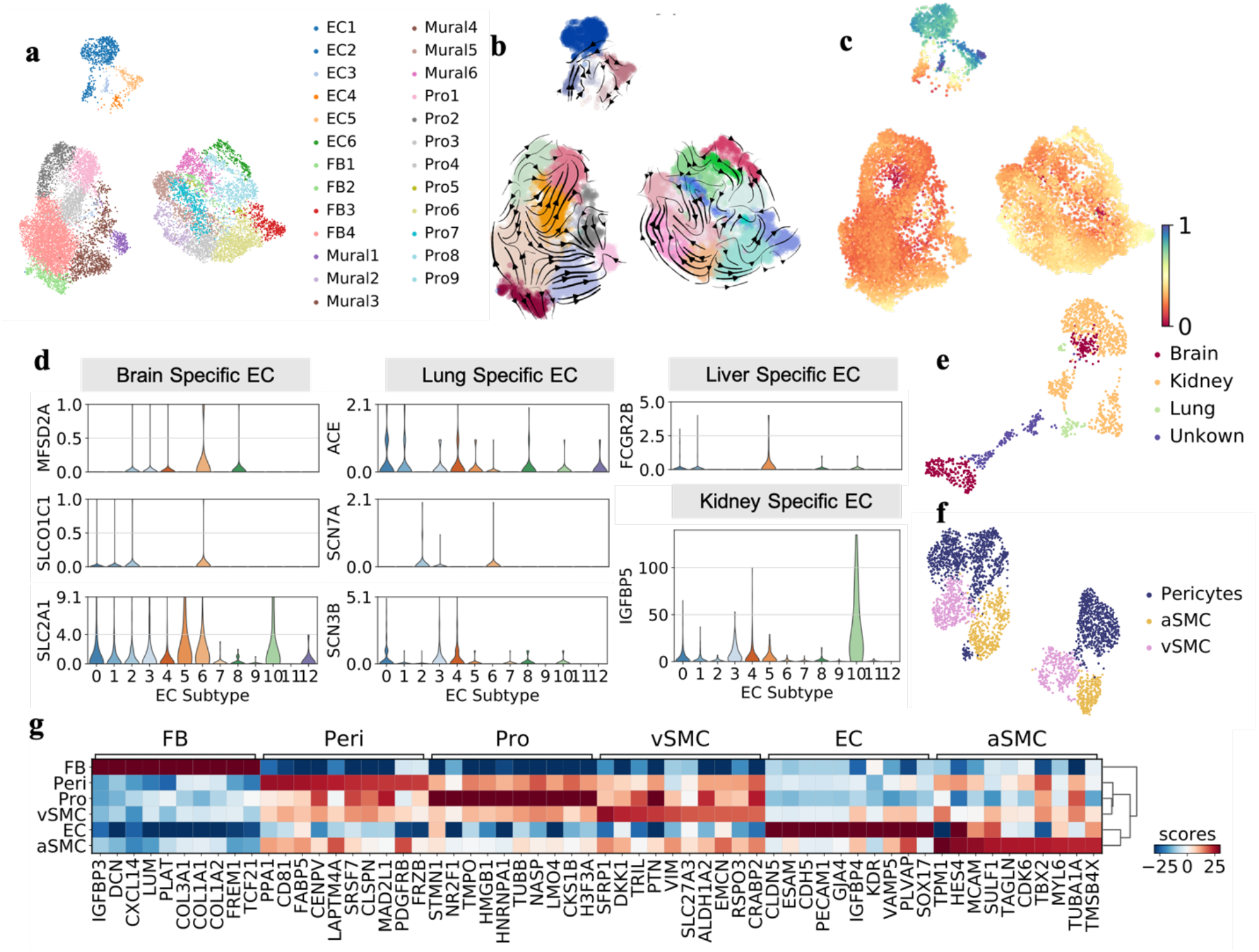
Single cell RNA profiling of blood vessel organoids. a) UMAP of sequenced cells in 22q11.2 DS and control VOs. Cell types have been determined by marker genes. b) Streamline plot of RNA velocity of case and control VO. The velocity indicated the developmental processes from progenitor to mature vascular associated cells. c) Pseudotime of the cells in the fig a) and b), the calculation was performed on control and case VO separately. Progenitors has shown as the cells at the earliest time point while the endothelial cells were at the latest time points. d) Organ specific gene expression in endothelial cells. e, f) UMAP of EC subtypes. g) Differential genes cluster map for cell types in VO.

### Variation of cell components and fate determination during 22q11.2 DS VO differentiation

Based on the characterization shown above, we explored the variation in the cell fate determination and developmental processes in the 22q11.2 DS VOs. Similar to the findings on morphology and topology characteristics that the disease VOs maintained the fundamental vascular structure and function, all the identified vascular cell types existed in both 22q11.2 DS and the control VOs. However, the percentages of EC and fibroblast (FB) were higher than those in the control VOs (EC: ∼11.1 % in 22q11 vs. ∼8 % in control; FB: ∼43.6 % vs. ∼10 %), while the percentages of mural and proliferating cells were less (mural: ∼15.8 % vs ∼29 %; proliferating: ∼29.6 % vs. ∼53 %; Fig. 6b). This is consistent with our suggestion of a high percentage of progenitor cells in the control (Fig. 1f). The comparison of EC subtypes demonstrated that the percentages of arterial EC were similar between the 22q11.2 DS and the control VOs, while the venous EC numbers were reduced, and capillary EC was significantly enriched. This finding suggests that the observation of thinner vascular tubes in the 22q11.2 DS VO from morphology analysis might be due to the enrichment of capillary (Fig. 6c). The organ specificity comparison demonstrated the developments of brain, kidney, lung-specific EC in both normal and disease VOs. However, the percent of kidney-related lineages was relatively higher, and brain-related lineages lower in 22q11.2 DS VO. As the 22q11.2 DS has been reported to affect both brain and kidney in clinical studies^43^, we focused on these two organ-specific EC types, respectively. We identified 570 differential genes in the brain EC (388 genes upregulated and 182 downregulated), and 131 differential genes in the kidney EC (64 upregulated and 68 downregulated; *p* <0.01, logFC >0.5; Fig. S5a). There were 78 genes that overlapped between brain- and kidney-specific ECs (Fig. S5b). Gene ontology analysis revealed downregulated genes in the 22q11.2 DS brain-specific EC are involved in cell division, proliferation, consistent with our morphological findings (Fig. 1f, Fig. 2g). Meanwhile, the downregulated biological processes involved in brain development and kidney development were consistent with the clinical phenotype^43^(Fig. S5c). The upregulated genes in the 22q11.2 DS brain-specific EC were associated with vasculogenesis, angiogenesis, endothelium development, and cell adhesion (Fig. S5c). The gene ontology analysis of downregulated genes in the kidney-specific EC showed the enrichment of the biological processes that were involved in smooth muscle cell development, angiogenesis sprouting, wound healing response, and paracrine signaling, while upregulated genes were associated with immune response, positive regulation of vasoconstriction, extracellular matrix organization, negative regulation of endothelial cell proliferation and collagen fibril organization (Fig. 6d). The cellular components of both upregulation and down-regulated genes were associated with focal adhesion, cell junction, and extracellular matrix. KEGG also demonstrated the effects on focal adhesion, filopodium membrane, ECM-receptor interaction, and *PI3K-AKT* signaling pathway (Fig. 6d). Collectively, we identified different cell-ECM and cell-cell contact of ECs in the 22q11.2 DS VO compared with the wild-type counterpart.

**Figure 6.**
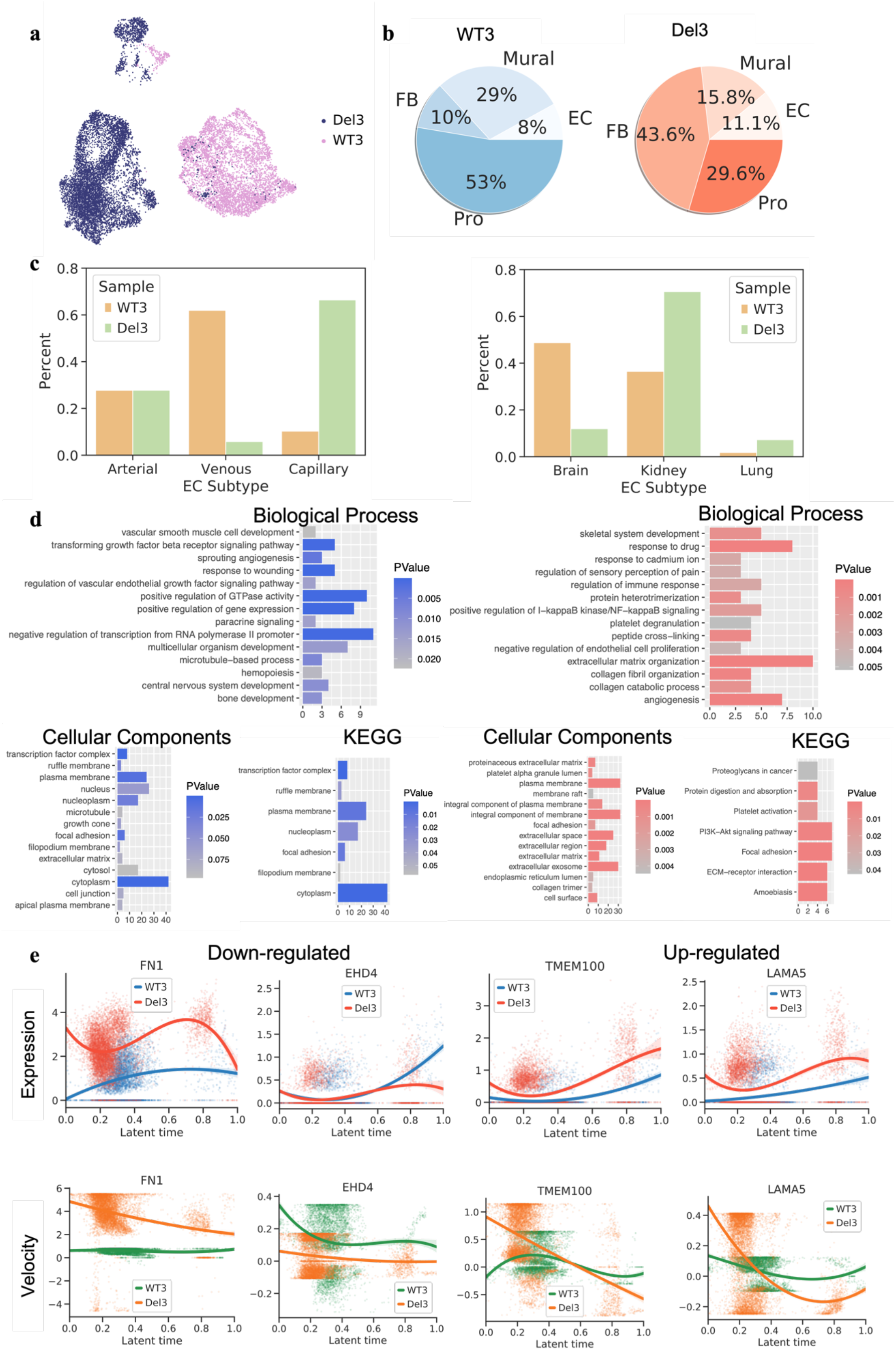
Variation in cell components and developmental processes in 22q11.2 DS VO. a) UMAP of sequenced cells in 22q11.2 DS and control VOs colored by sample (Del3: 22q11.2 DS VO, WT3: control VO). b) Pie plot of cell types in 22q11.2DS VO and control VO. c) Comparison of EC subtype and organ specificity in 22q11.2 DS VO and control. d) Gene ontology analysis including biological processes, cellular components, and KEGG signaling pathways of downregulated and upregulated genes for kidney like EC in the 22q11.2 DS VO compared to control VO. e) RNA expression and velocity along with the latent time of representative variable genes in ontology analysis in d).

To understand the differences in lineage dynamics between the 22q11.2 DS and control VOs, we then investigated the RNA velocity change along with the pseudotime. Fig S5d displayed two heatmaps for RNA velocity change along with latent time in the disease and the control VOs, which indicated the expression fluctuation of those variable genes that determined the developmental processes. We explored those marker genes for the cell type determination. As shown in Fig S5e, the expression and velocity of PECAM1 indicated the maturation of EC in 22q11.2 DS was earlier than that in the control, The velocity of PECAM1 indicated some cells had active transcription processes at around 0.2 (EC progenitor) and 0.8 (mature EC) of the latent time.

There was a large decrease of velocity to the negative after 0.8, indicating the maturity of some EC cells in the disease VOs. The expression of PDGFRβ in 22q11.2 DS representing the mural cells was also dominant at an earlier time compared with the control. For both 22q11.2 and control, the maturation of mural cells happened around the same time (0.2 of the latent time). We also noticed the positive RNA velocity of PDGFRβ was ahead of PECAM1, suggesting a supportive role of mural cells for EC development. Then we investigated the genes involved in the identified KEGG pathway and biological processes. FN1 was the gene associated with ECM-receptor interactions, cell adhesion, wound healing, and focal adhesion, where we saw its higher expression and velocity among all the latent time, implying the enhanced ECM-receptor interactions in the 22q11.2 DS VO. EHD4 was involved in cell-cell adhesion, where we saw lower RNA velocity in the disease VO over the control. In addition, the higher velocity of TMEM100 (involved in angiogenesis) revealed the enrichment of vascular development in our disease VO. We also noticed a higher expression level of LAMA5 in the 22q11.2 DS VO, a well-known regulator in basal lamina and extracellular matrix, indicating the enrichment of ECM in the 22q11.2 DS VO. Overall, the variation in the cell components revealed the dynamical regulation of hub genes that led to the dysfunction in the 22q11.2 DS vasculature.

### Alterations in the intercellular communication and functional signaling pathways in 22q11.2DS VOs

To understand the mechanism underlying the affected biological processes during the vasculogenic developments in the 22q11.2DS VOs (e.g., how different cell type changed their role in those signaling pathways), we investigated the intercellular communications from our scRNA-seq data, through the analysis of the signaling pathways associated with those critical variable genes. By implementing CellChat^44^, a method for inferring the intercellular communications by combining network analysis, pattern recognition, and manifold learning, we first identified the global alterations regarding the number of inferred molecular interactions and strength. We noticed that the 22q11.2 DS VOs exhibited a lower number and strength of significant molecular integrations associated with cell-cell contact (149 in 22q11.2 DS vs. 296 in the control for the number of cell-cell contact and 5.946 vs. 6.079 for the strength of cell-cell contact; Fig. 7a). In contrast, the number and strength of molecular interactions associated with the extracellular matrix (ECM)-receptor and secreted signaling in the 22q11.2 DS were significantly higher (ECM Receptor: 641 vs. 364 for the number and 39.029 vs. 7.891 for the strength; secreted signaling: 238 vs. 195 for the number and 4.702 vs. 2.075 for strength). We then classified the data into 17 signaling pathways related to cell-cell contacts (e.g., CDH5, NCAM, NOTCH, PECAM1, PTPRM), 5 signaling pathways related to ECM-receptor (COLLAGEN, FN1, LAMININ, TENASCIN, THBS), and 19 secreted signaling pathway (e.g., FGF, PDGF, VEGF, IGF, PTN). The relative information flow in each signaling between 22q11.2DS VO and control is summarized in Fig. 7b. We then explored the differential number of interactions and strength in different cell types by network centrality analysis (Fig. S6a). The number of cell-cell contact signalings was consistently decreased in many cell types in 22q11.2 DS VO, although the interactions between pericytes, SMC, and proliferating cells had a small increase. The cell-cell contact strength showed an increased information flow from all other cell types to ECs, while ECs sent fewer signals to all other cell types in 22q11.2 DS VOs. The number and strength of ECM-receptor signalings in the 22q11.2 DS VOs were increased between all the cell types except the signals from SMC to itself and from SMC to pericytes, which is decreased. We also found the number and strength of secreted signals were highly increased from mural cells, fibroblasts, proliferating cells to EC, while the number of secreted signaling from EC to other cell types decreased. These findings indicated that the ECs have a deficiency in sending signals, especially for cell-cell contact. Next, we explored the major sources and targets among cell types. The scatterplot (Fig. 7c) presents the incoming and outgoing interaction number and strength for different cell types between the 22q11.2DS and control VOs in three types of signaling pathways. We found the enhancement of incoming interaction of cell-cell contact (increased from ∼1.4 to ∼2.4) and secreted signaling (increased from ∼0.4 to ∼1.7), as well as the outgoing secretion signals (increased from ∼0.26 to 1.00) of ECs. However, the outgoing interactions of cell-cell contact of ECs reduced from ∼1.6 to ∼1.3, which led to the reduction of incoming interactions in other cell types. We also identified that the SMC and pericytes were the two major cell types with the most enriched ECM-receptor signaling including incoming and outgoing. We thus hypothesized that the vasculopathy in the 22q11.2 DS was due to the weakness of EC as a signal sender, inducing an increase in ECM-mural cell interaction, such as focal adhesion. To confirm our hypothesize, we investigated the contribution of individual signaling pathways in each cell type. By joint manifold learning and classification of inferred communication networks based on the topological similarity, CellChat was able to identify the most altered signaling pathways (Fig. S6b), where we identified CD99, collagen, and VEGF as the most altered cell-contact, ECM-receptor and secreted signaling pathways. This finding also suggested that the vascular formation and ECM remodeling were the most important processes in developing VOs. As a result, we analyzed the signaling role of VEGF, collagen, and CD99. We found that EC was the main cell type for sending, receiving, and influencing in the VEGF signaling pathway, and SMC acted as an influencer as well. In the control VO, EC was the only sender and receiver of VEGF signaling and there was no other mediator or influencer found (Fig. S6c, Fig. 7d). In contrast, in the 22q11.2 DS VO, other cells, including FB and pericytes, played roles as sender, mediator and influencer in the VEGF signaling, to target EC. We also identified that FB was the main sender and SMC was the main receiver of COLLAGEN signaling in control VOs; However, the 22q11.2 DS pericytes played a more important role in pericytes than FB including all four roles. These results are also consistent with the single-cell gene ontology analysis. Bulk RNA analysis of another 22q11.2DS pair (Del2/WT2) also showed upregulation of extracellular matrix organization and angiogenesis (Fig. S7). Our results showed the significance of EC and SMC in the CD99 signaling was lost and replaced by FB. This finding is consistent with weaker colocalization of EC and mural cells in 22q11.2 DS VO. Overall, the investigation of intercellular communication and gene ontology analysis suggested that the morphological changes in 22q11.2 DS VO might be caused by inter-cellular communication problems.

**Figure 7.**
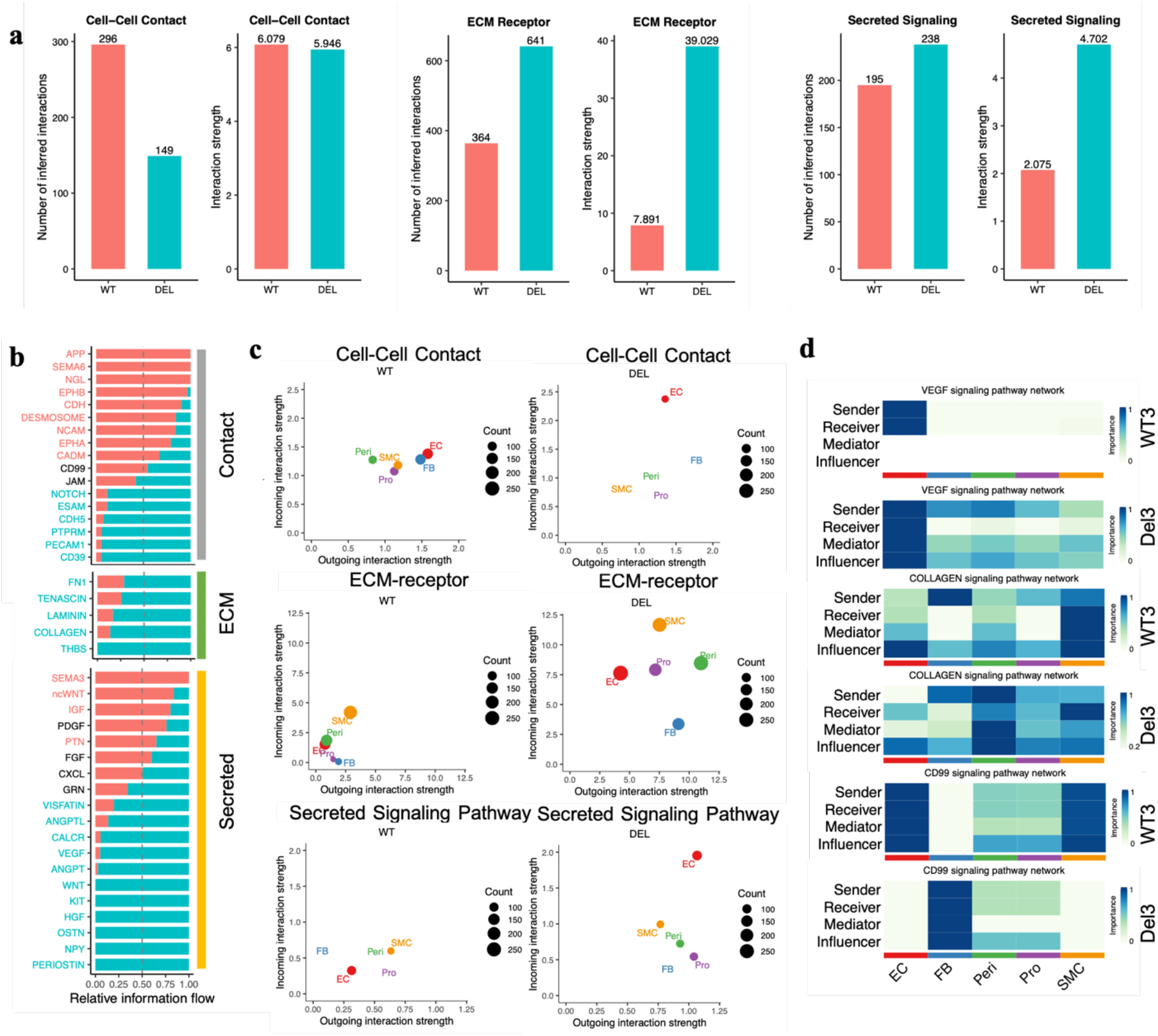
Variation in the intercellular communications and functional signal pathways in 22q11.2 DS VO. a) Number of inferred interactions and strength in cell-cell contact, ECM receptor, and secreted signaling pathways. b) Identified variable signaling pathways. c) Relationship between incoming and outgoing interaction strength. d) Significant signaling identified from WT and DEL. d) Roles of different cells in VEGF, COLLAGEN, and CD99 signaling pathways.

## Discussion

The iPSC-derived VO models the patient-specific vascular system by recapitulating the specific vascular variation caused by the disease. It can thus act as a transformative *in vitro* biomimetic platform for understanding genetic lesion-related vasculopathy and analyzing vascular diseases beyond the 22q11.2 DS model presented in this work. Here, we leveraged the integration of deep learning-based segmentation, network graph construction, tessellation analysis, and single-cell RNA profiling for characterizing vascular structures and cell compositions, understanding the genetic function, and enabling a comprehensive way to illuminate the pathogenesis of vasculopathy in 22q11.2 deletion from a new angle. The 22q11.2 DS VOs reveal nuanced vascular morphological differences compared to their wild-type counterparts. The smaller VO size with 22q11.2 deletion and GO analysis pointed out impaired cell proliferation in the disease. The smaller vascular diameters of endothelial tubes indicated an abnormality of tubular formation, which may be due to the immaturity of the smooth muscle cells. The smaller tubular diameters also may be due to the enriched capillary identified from single-cell RNAseq. Quantification of network connectivity of VO elucidated the complexity of the vascular network and efficiency of blood flow in the vascular systems. Although there was enhanced vascular density, angiogenic sprouting, and enriched bifurcation points in the VOs (and in other studies^6^), the vascular network connectivity was significantly lower, which may lead to an ineffective interchange of material. Further network investigation considered the VO as an individual network graph and the degree exponent was used to demonstrate the lack of robustness in 22q11.2 DS VO and its susceptibility to the environmental perturbation, which underlies the deficiency of the vascular system due to 22q11.2 deletion. The mechanism of the pathogenesis of 22q11.2 deletion vasculopathy was further revealed through the combination of morphology analysis and single-cell RNA sequencing. The characterization of single-cell atlas for vessel organoids constructed through the analysis of single-cell transcriptome enables the comparison of VO heterogeneity and development between 22q11 deletion and control. Cell type composition in our generated VOs included EC, vascular associated cells (pericytes, smooth muscle cells, and fibroblast). Exploration of EC subtypes including organ specificity and GO analysis revealed a kidney development deficiency in the 22q11.2 DS, consistent with the clinical diagnosis^43^. The decreased cell-cell contact identified through the GO analysis and intercellular communication analysis matches the indication of less direct interaction between EC and mural cells through the colocalization analysis. Other affected biological processes and pathways included the abnormality and cell dysfunction in cell adhesion, PI3K-AKT-VEGFR signaling pathways, ECM-receptor, and ECM remodeling in VO with 22q11.2 deletion. The consistent findings from morphological structure analysis, colocalization study, and transcriptome profiling of the vessel organoids all demonstrate the value of an integration of stem cell engineering and machine learning for modeling complex vascular diseases and potentially preclinical drug screening.

## Methods

### Reprogramming and culture of human pluripotent stem cells (iPSCs)

Human iPSC lines were generated from three patients with 22q11.2DS and three healthy controls as described previously^45^. Human fibroblast and blood cells were reprogrammed with a Sendai virus as previously described^45^. Karyotyping and genome sequencing were performed for quality control and genotype validation.

The iPSCs were cultured in mTeSR plus medium (STEMCELL, 05825) with Matrigel (Corning, 354277) coated 6-well plates. Gentamicin (Gibco) with 50ug/ml was added as an antibiotic. The media was changed every day, and cells were passaged with Accutase (Thermo Fisher, A1110501; Sigma A6964) every 3-4 days.

### Generation of blood vessel organoids

Vessel organoids were generated with 6 hiPSC lines using a previously published protocol, ^15^ with minor modifications. Briefly, hiPSCs were dissociated with Accutase and triturated to generate single-cell suspensions. A total of 277000 cells were plated into one well of ultra-low attachment Elpasia 24-well plate (Corning, 4441) to form embryoid bodies (EBs) with mTeSR plus medium containing 50 μM chroman 1 (MedCHemExpress, HY-15392), 1.5 μM emricasan (Selleck Chemical, S7775), polyamines (Sigma Aldrich, P8483), and 0.7 μM trans-ISRIB (Tocris Bioscience, 5284) for the first 24 hours ^46^. The medium was switched to N2B27 medium containing 50% DMEM/F12 (1:1) (Gibco; 12660012), 50% NeuroBasal (Gibco, 21103049), GlutaMAX (Gibco, 35050061), N-2 supplement (Gibco, 17502048), B27 supplement minus vitamin A (Gibco, 12587010), Gentamicin (Gibco, 15710064), and 2-Mercaptoethanol (Gibco, 21985023) with the addition of 12 μM CHIR99021 and 30 ng/mL BMP4 for the following three days. On day 3, the medium was switched to N2B27 medium supplemented with 100 ng/mL VEGFA (ProSpec, CYT-116) and 2 μM forskolin (Tocris Bioscience, 1099) for two days. On day 5, the spheroids were embedded in a Matrigel-collagen I (Corning, 40236) mix (1:4) in medium containing DMEM/F12, 15% FBS, 20% knockout serum replacement (Gibco, 10828), GlutaMAX, MEM-NEAA (Gibco, 11140050), and 2-Mercaptoethanol with addition of 100 ng/mL VEGFA and 100 ng/mL FGF2 (ProSpec, CYT557) for 4 days. Blood vessel sprouting was observed from day 7 and was well-developed at day 10. Those vessel organoids were then fixed with 4% paraformaldehyde at day 12 for imaging or dissociated for single-cell RNA or bulk RNA sequencing. All iPSCs and organoids were maintained under well-defined conditions avoiding mycoplasma contamination.

### Whole-mount Immunocytochemistry

Vessel organoids (day 11) were fixed with 4% paraformaldehyde (PFA) overnight at 4°C followed by three times of PBS was for one hour. Samples were blocked with blocking buffer (PBS with 3% FBS, 1% BSA, 0.5% Triton X-100 and 0.5% Tween-20) at 4°C overnight. To preserve the fine structure of the vascular network, organoids were transferred together with the gel and placed into 50 ml tubes. Primary antibodies were diluted at 1:100-1:200 ratio in blocking buffer, and organoids were incubated in primary antibodies overnight at 4°C. The following primary antibodies were used: anti-human CD31 (Abcam, ab28364) and anti-human PDGFR-β (R&D Systems, AF385). After three 8-hour washes in PBST at 4°C, samples were incubated with the corresponding secondary antibodies overnight at 4°C: Alexa Fluor 647 donkey anti-goat (Jackson Immuno Research Labs, 705606147) and Alexa Fluor 488 donkey anti-rabbit (Jackson Immuno Research Labs, 711546152). After three 8-hour washes in TBST at 4°C, samples were counterstained with DAPI overnight at 4°C followed by a 4-hour wash in TBST.

### Lightsheet microscopy

Stained vessel organoids were transferred to 50 mL tubes. The solution residue was removed prior to tissue clearing with RapiClear (SunJin Lab, RC149001) at room temperature until reaching the homogenous refractive index 1.49nD. Cleared samples were mounted in a 4-window quartz cuvette and imaged by a light-sheet microscope, as previously described^47^. The confocal microscope 10X magnification was chosen for imaging. Images achieved 0.585um per voxel on the x and y axes and 5um on the z direction. Imaging tiles were stitched before performing the analysis. Organoids derived with the same cell lines were imaged together embedding in the oil in one quartz cuvette.

### Imaging pre-processing

Region of interests (ROI) with individual blood vessel organoids were captured and saved as separate files. To increase the quality of images, homomorphic filtering and contrast limited adaptive histogram equalization were applied to the original datasets with the same parameters. Normalization of imaging intensities was applied to adjust the intensities to the range of 0-1.

### Deep learning of vessel organoids segmentation

To segment the CD31+ endothelial tubes, we adopted the 3D convolutional neural network (CNN) model from VesSAP^21^. The model contained five convolutional neural network layers, each with full 3D convolutional kernels and ReLU activation function. The last layer was a fully connected layer and followed by Sigmoid activation to project the prediction of segmentation into the range of 0-1. A threshold value of 0.9 was set to determine the segmentation from generated probability maps. We used the trained parameters on whole mouse brain vasculature^14^ and simulated vascular trees^14^ due to the scarcity of training data, and similar data features between mouse brain capillary and blood vessel organoids^14^, ^48^. The data input of the model had two channels; thus, we duplicated the CD31+ channels. The results were able to capture the important network features such as branch junction and 3D direction of each vascular branch.

The segmentation of nuclei in vessel organoids was implemented by the Cellpose^23^, a deep learning-based method capable of segmenting DAPI stained nuclei in the 3D space. To achieve high accuracy in 3D space, we first estimated the nuclei diameters in 2D space and then used the results for parameters initialization in the 3D prediction.

We then implemented the ilastik^49^ to perform 2D segmentation on PDGFRb+ mural cells. This data didn’t harbor a well-formed network structure but connected to a certain shape instead of separated nuclei distribution. Ilastik provides a user interface on training processes that provided fast feedback in training, thus reducing the workload and performing fast segmentation tasks.

### Morphological and topological analysis

Skeletonization was performed based on the results of vascular segmentation. The preprocessing for skeletonization included binarization, binary closing, and filling holes to correct false-negative pixels. Skeletonization was performed by using the function ‘skeletonize_3d’ in the python library ‘skimage’, which is based on the 3D thinning algorithm. Skeleton was filtered by removing small objects. Gaussian kernel density estimation of the skeleton was performed to estimate the probability density function (PDF) of the vascular tubes. Two Python libraries, Skan and Networkx, were utilized for the analysis of skeleton and network graphs. Bifurcation points were identified based on the skeleton of vascular tubes and classified into three types of points: end points, path points, and joint points, as explained in Fig. 2b. Graph *G* = {*N, E*} was created where *x*_*i*_ ∈ *N* represents bifurcation points and *e*_*i,j*_∈ *E* denotes whether *x*_*i*_ and *x*_*j*_ was connected directly. The degree of a node was defined as the number of connections it had with other nodes. The touristy was calculated by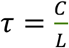, where *c* denoted to the length of the curved branch length, *L* denoted to the Euclidian distance between the two ends of the branch.

## Colocalization analysis

The colocalization analysis was based on the Voronoi diagram and the previously published method^31^, ^50^. The Voronoi diagram partitioned the space of vessel organoids containing an ensemble of binarized signals into small polytopes. The binarized signals included pixels with segmentation probability >0.9 in endothelial network, or >0.5 in mural cells. Their locations were denoted as 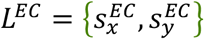 and 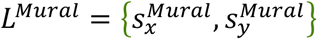, and their corresponding voronoi polytopes were denoted as *P*^*EC*^ and *P*^*Mural*^, the area of each polytopes were calculated and denoted as *A*^*EC*^ and *A*^*Mural*^. The 1^st^ rank local localization density *δ*_*i*_ was computed as: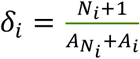, where *N*_*i*_ was the number of neighbor spots those had adjacent voronoi polytopes, *A*_*i*_ was the area of polytope of spot 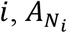 was the area summation of adjacent voronoi polytopes. *δ*_*i*_ was normalized by 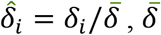 referred to the average value in the space. Each voronoi region was classified into high density and background based on the threshold, which was set to T=1 in this study. The processes could be denoted: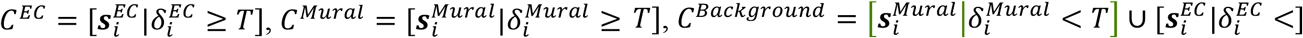 . Adapted the method published in ^31^, we used pair density descriptor 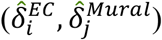 and 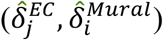 to quantify the spatial correlation of endothelial tubes and mural cells, where 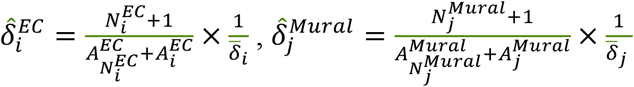 and *i* and *j* had overlapped voronoi diagram. Thus, the classes could be updated as 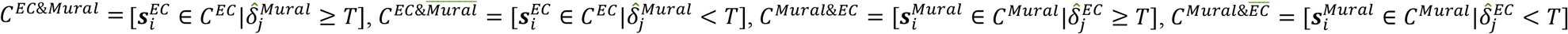 . The voronoi manders’ overlapping coefficients were defined as: 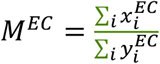, where 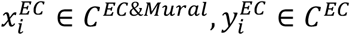 and 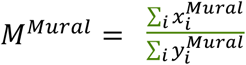, where 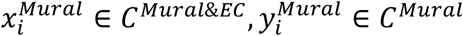 . Spearman’s rank correlation coefficients was calculates as 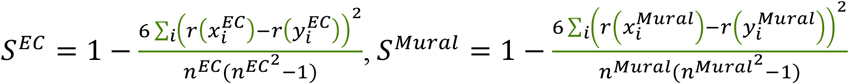, where *r*(*x*_*i*_), *r*(*y*_*i*_) was the ranks of each signal spots in EC and mural cell channels.

### Single-cell RNA sequencing and analysis

Based on the morphological and topological analysis, VOs generated from WT3 and Del3 iPSCs lines have displayed the most significant differences in the morphology. Thus WT3 and Del3 VOs (at Day 11) were gently and completely dissociated into single cells using Accutase. Cell viability was confirmed as 90% and 87% in two samples by cell counter and trypan blue to ensure the reliability of the subsequent sequencing results. Cells were washed three times with PBS, and suspended single cells were captured by barcoded beads using 10X Genomics Chromium technology. Cellranger v3.1.0 was performed on the raw sequencing data. The generated raw unique molecular identifier (UMI) count matrix was then processed by Scanpy ^51^. Cells with UMI counts less than 7000 were filtered out, and with 10% mitochondrial gene were also removed. And we also filtered out the cells expressing less than 1200 genes, and genes with expression in at less than 3 cells. Raw UMI counts were then normalized by the median of the library size, and log transformed. Then we used scanpy to identify 10000 highly variable genes for the following analysis. Principle component analysis then was performed (PCA) and the number of PC was determined through the cumulative distribution function, which turned out as 83 PCs. Then Phenograph was performed on the top 83 PCs. UMAP was also performed for purpose of visualization. To annotate the cell types in each cluster, we relied on well-studied marker genes^14^ for each vascular associated cell type to confirm the cell types (endothelial cell: PECAM1, CLDN5, FLT1, CDH5; pericyte: PDGFRb, AMBP, DES, VTN; smooth muscle cells: ATAC2, TAGLN; Fibroblast: COL1A1, CAL1A2, LUM). We also performed gene set enrichment to confirm the cell type annotation. RNA velocity analysis was performed based on the scVelo^39^, ^40^. RNA velocity of cells in 22q11.2 and control VO were calculated separately and visualized on the same map. Pseudo-time of each cell was obtained from the RNA velocity analysis. Cell-cell communication analysis was performed by the method Cellchat^44^.

### Bulk RNA Preparation and analysis

More than 20 vessel organoids per line were collected at day 11. Total RNA was extracted using Direct-zol RNA Miniprep kit (Zymo Research, R2052) according to the instructions of the manufacturer. RNA eluted with RNase-free water was assayed by spectrophotometer (Nanodrop) to determine the purity and concentration. Extracted RNA was reverse transcribed into cDNA with iScript cDNA synthesis kit (BioRad, 1708891). Bulk RNA samples were purified from ∼30 vessel organoids derived from each cell line in 5mL of Trizol and standard Qiaquick RNA extraction protocols. Stranded RNA-seq libraries were generated at the Columbia Genome Center using a polyA pulldown method. Stranded cDNA libraries were paired-end sequenced (2×125 bps) on a HiSeq2500, operating in high output mode and yielding 30M reads per indexed library.

The reads were pseudo-aligned with the R package Kalisto and mapped to the human genome. The matrix was further analyzed with the web-based tool iDEP^52^. In brief, a significance threshold of FDR <0.1 and log2 fold change>2 was applied to filter the differentially expressed genes with the embedded DEseq2 package.

### Statistics

Statistical analyses were performed using GraphPad Prism, Python, and Matlab. For comparisons of vascular structures between the controls and cases, we used Mann-Whitney tests, with a p-value of <0.05 considered as statistically significant. Data were presented as mean ± s.e.m. Statistical tests and biological replicates for each experiment were presented in the figure legends.

## Author Contributions (TBD)

S.H., C.X., and K.W.L. conceived the project, S.H., C.X. performed the organoids generation, S.C. performed the COLM imaging, C.X. did bulk RNA-seq preparation, S.H. did scRNA-seq preparation, S.H., and C.X. carried out the analysis. E.A., B.X. and K.W.L. supervised the project. All authors discussed the results and wrote the manuscript.

## Notes

The authors declare no financial competing interests.

## ACKNOWLEDGMENT

This work was supported by NIH UH3TR002151 and the Columbia Facility Core. S.H. also thanks Xueer Chen for scientific discussion.

## Supplementary Materials

1. Reagent information.
2. Antibody information.
3. Video of blood vessel organoids reconstruction.
4. Genomic validation of hiPSCs
5. Karyotypes validation of the 7 hiPSCs
6. Primers for qPCR
7. Gene list

## Data and code availability

The data and code supporting the findings of this study are included in the main text. Supplementary information is available at https://github.com/siyuh/22q11.2VO. The single-cell RNA-seq data will be accessible on Gene Expression Omnibus (GEO).

---

**Figure S1.**
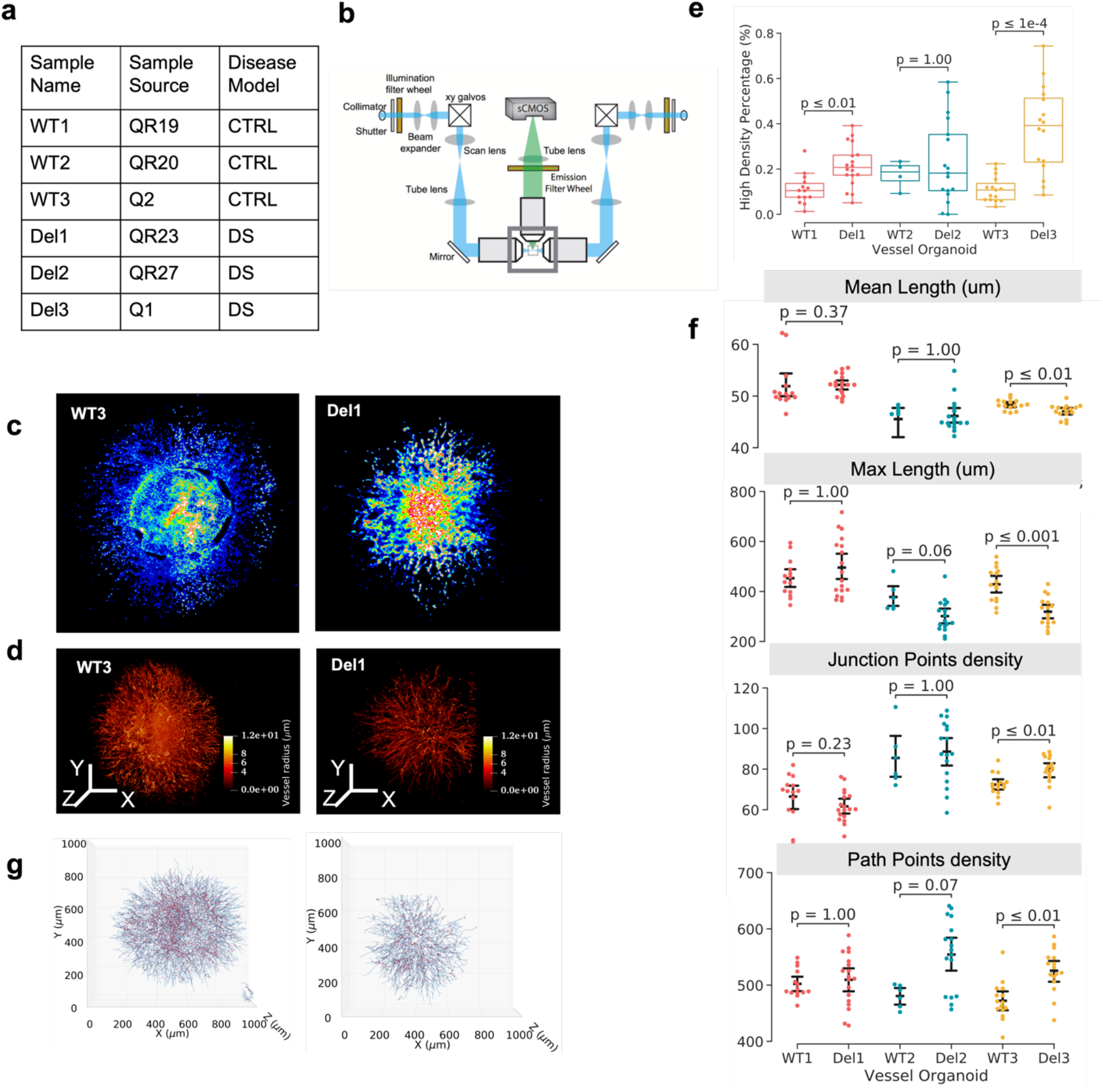
Light sheet imaging and morphological statistical summary of vascular l,k,k,structure in VO. a) Information of iPSC lines in this study. Three lines WT1, WT2, and WT3 were control. Del1, Del2, Del3 were corresponding 22q11.2 patient lines. b) Diagram of COLM. c) Heatmap of midplane of DAPI staining. d) 3D rendering of segmented control and 22q11.2 DS VO. e) Statistical comparison of high-density percent in VO (fold change = 0.9 and 2.4, *p* = 4.5×10^−3^ and 4.5×10^−5^ in WT1 and WT3, respectively; *n* = 84). f) Morphological feature of VO. (Mean length: Length_mean_ = 51.90, 45.53, 48.34 µm in WT1, WT2, WT3; Length_mean_ = 52.17, 46.13, 47.08 υm for DEL1, DEL2, DEL3, *p* = 0.37, 1.00, 9.70×10^−3^ respectively; fc = 0.005, 0.012, 0.026; Max length: Length_max_ = 453.00, 377.93, 428.59 µm for WT1, WT2, WT3, Length_max_ = 495.71, 301.12, 318.99 µm for DEL1, DEL2, DEL3, fc = 0.094, -0.203, -0.256; *n* = 84). e) Skeletonization of VO, the blue lines and red dots represent the centerlines of the microvascular tube and the bifurcation points, respectively. g) Skeletonization of VO.

**Figure S2.**
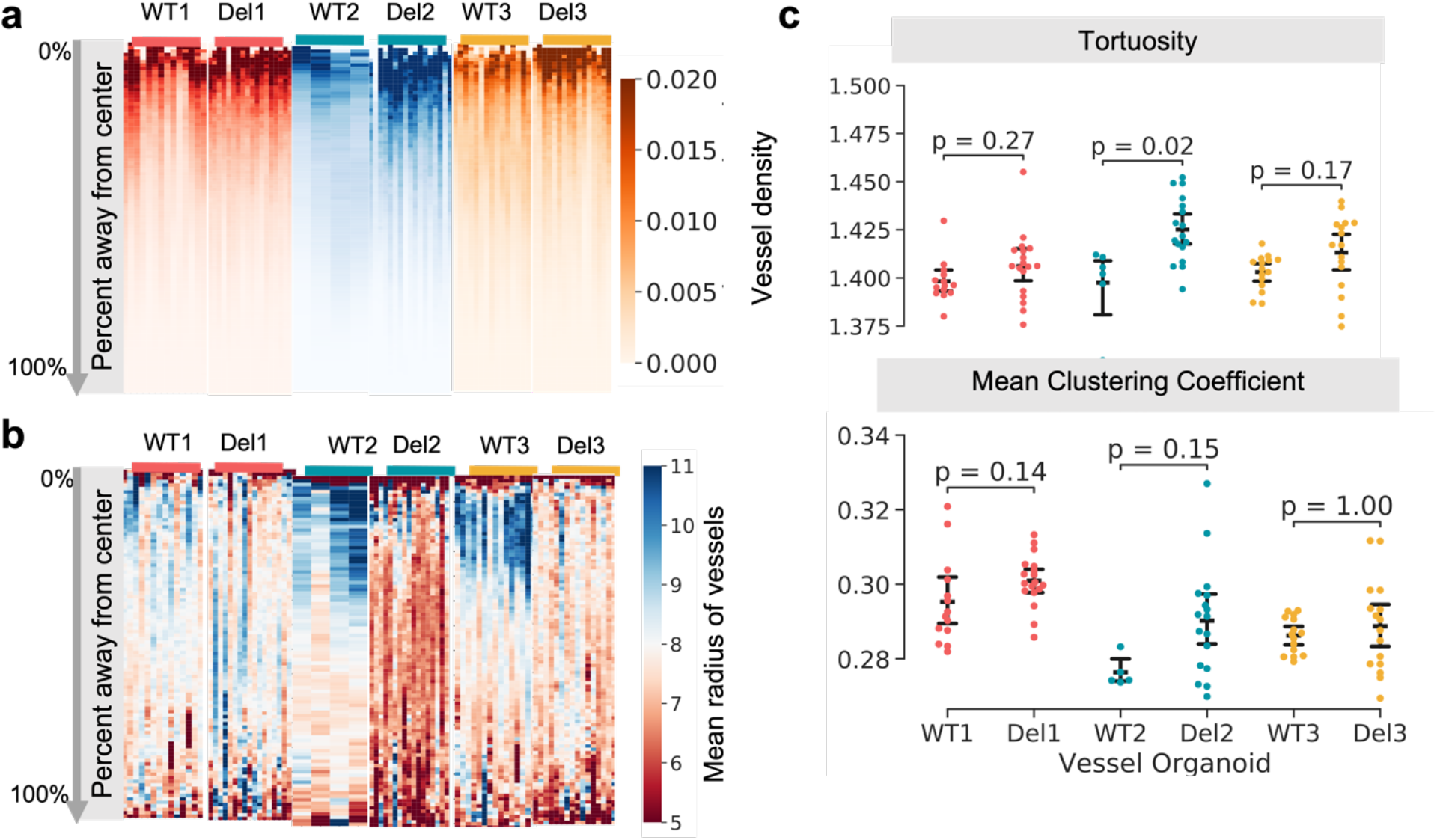
Morphological vascular features in radial coordinates. a) Heatmap of vascular density via the radial coordinates. b) Heatmap of mean radius via the radial coordinates. c) Tortuosity and clustering coefficient of VOs (Tortuosity: *p* = 2.728×10^−1^, 1.705×10^−2^, 1.657×10^−1^, clustering coefficient: *p* = 1.384×10^−1^, 1.505×10^−1^, 1.00).

**Figure S3.**
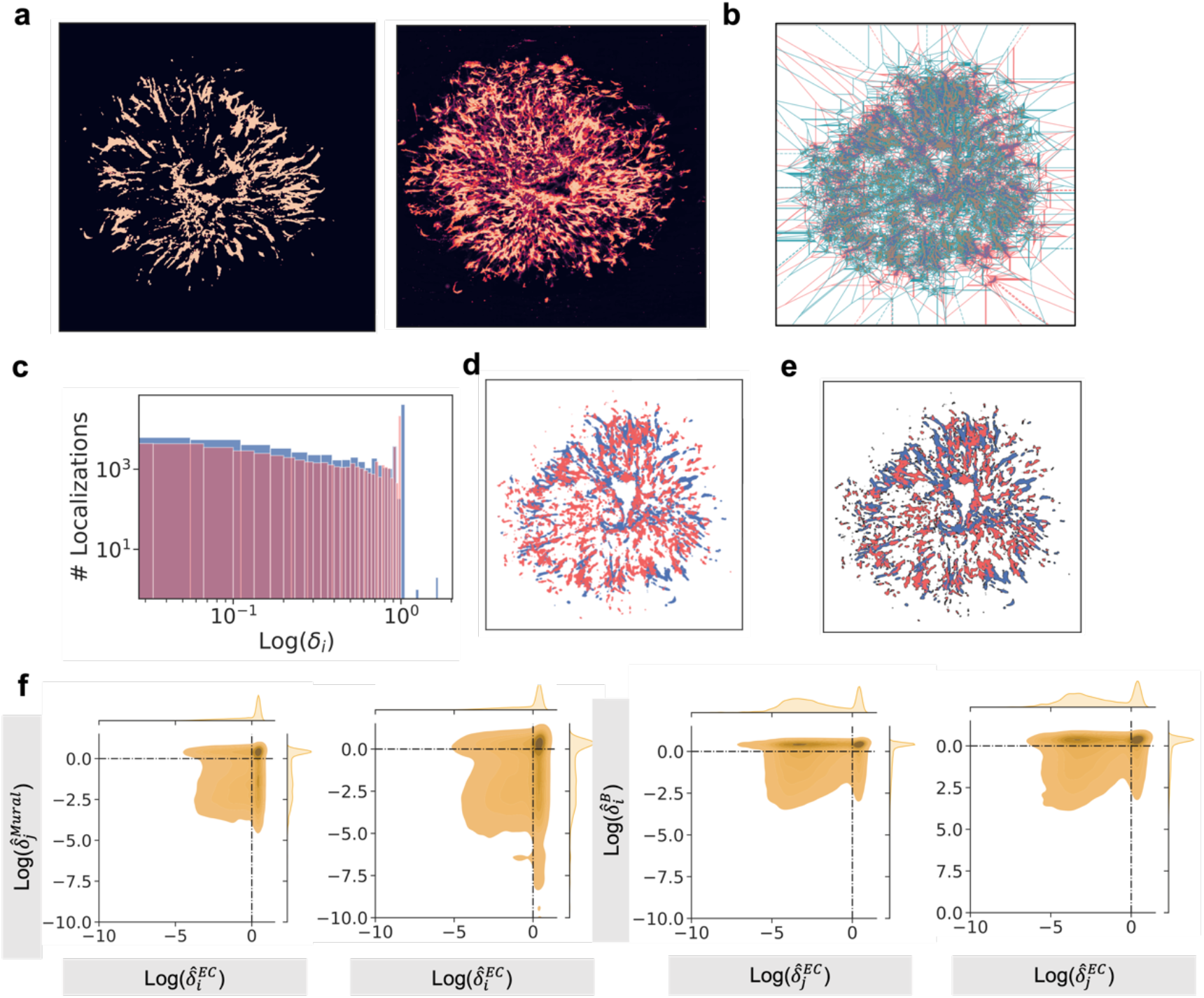
Colocalization analysis of vessel organoids. a) Segmentation of vascular tubes and mural cells by deep learning. b) Co-scatter plot of a). c) Distribution of normalized 1^st^ rank density. d) Voronoi diagram of two channels, indicating the overlap between two diagrams. e) Classification of the signals: high-density and background. f) Scatterplot of densities pairs of endothelial and mural cells in WT3 and Del3 VO.

**Figure S4.**
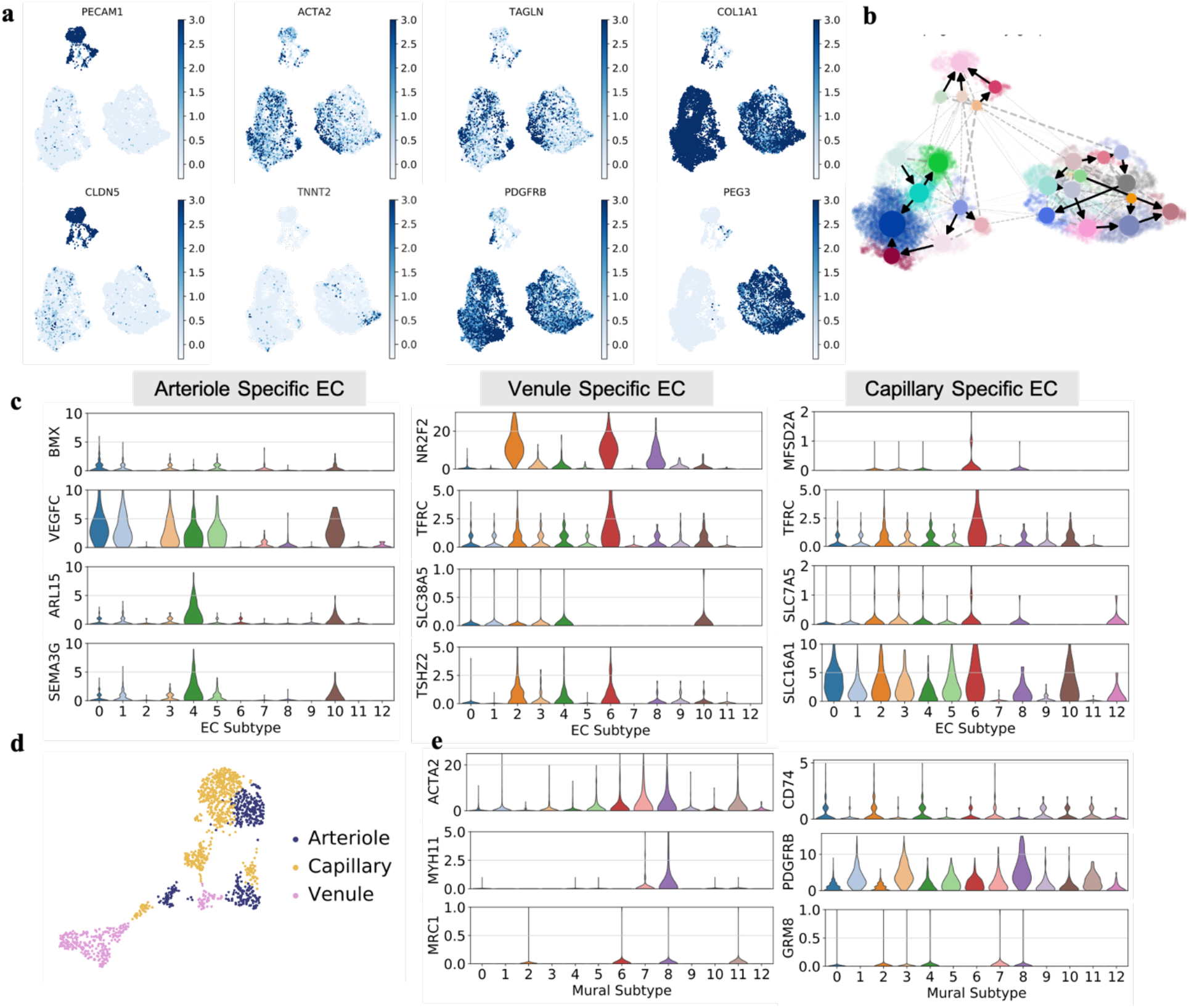
Heterogeneity of vessel organoids. a) UMAP with vascular associated marker genes. b) PAGA of cell clusters. C) Violin plots of EC subtype marker genes. d) UMAP of EC subtypes in all the sequenced ECs. e) Violin plots of Mural subtype marker genes.

**Figure S5.**
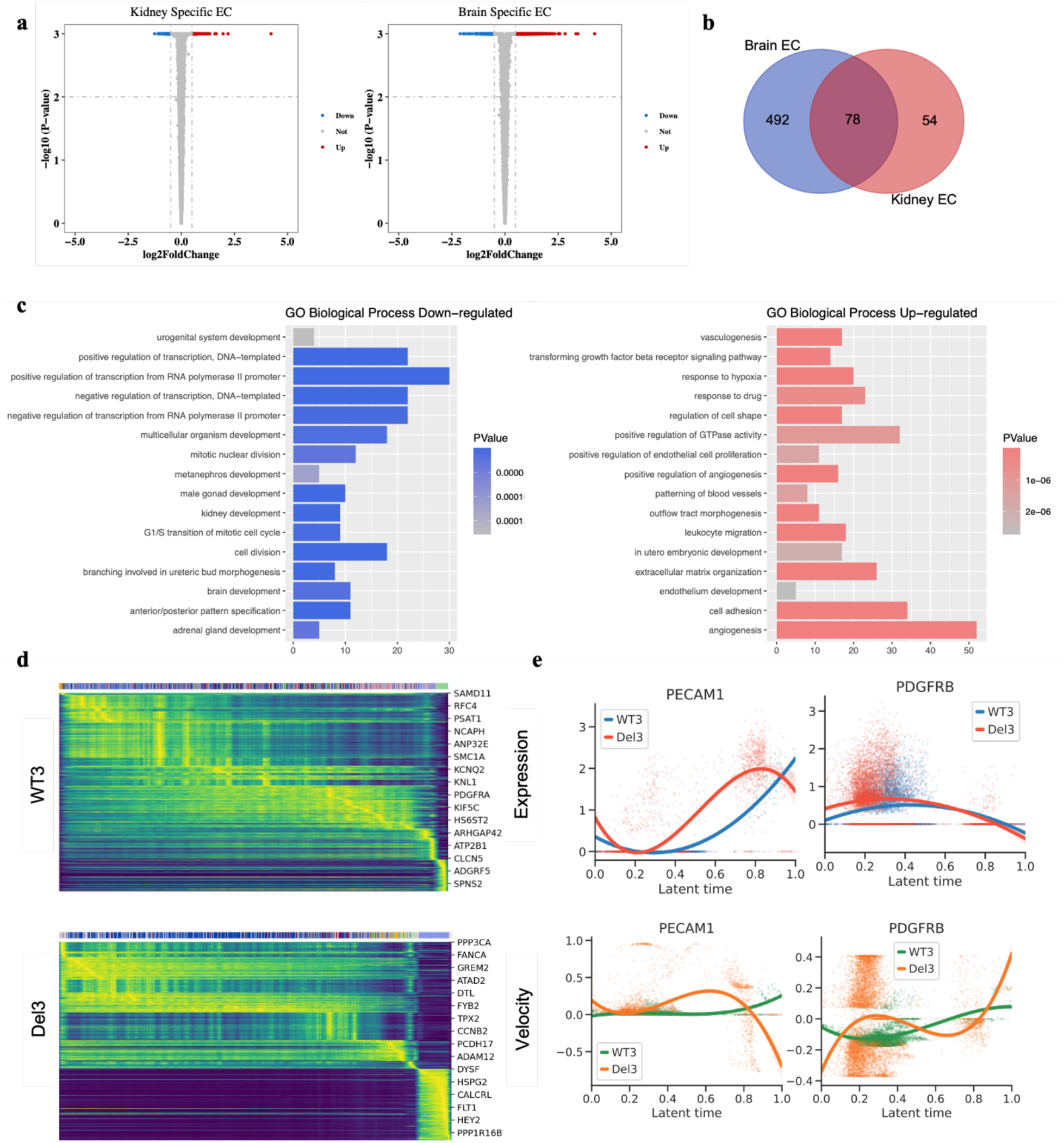
Heterogeneity of fate determination in 22q11.2DS vessel organoids. a) Volcano plot of differential genes in brain and kidney specific EC. B) Venn plot to show the shared differential genes in brain and kidney EC. c) Gene enrichment analysis in brain specific EC. d) Heatmap of genes among the latent time. e) RNA expression and velocity of EC and mural cell marker genes along with the latent time.

**Figure S6.**
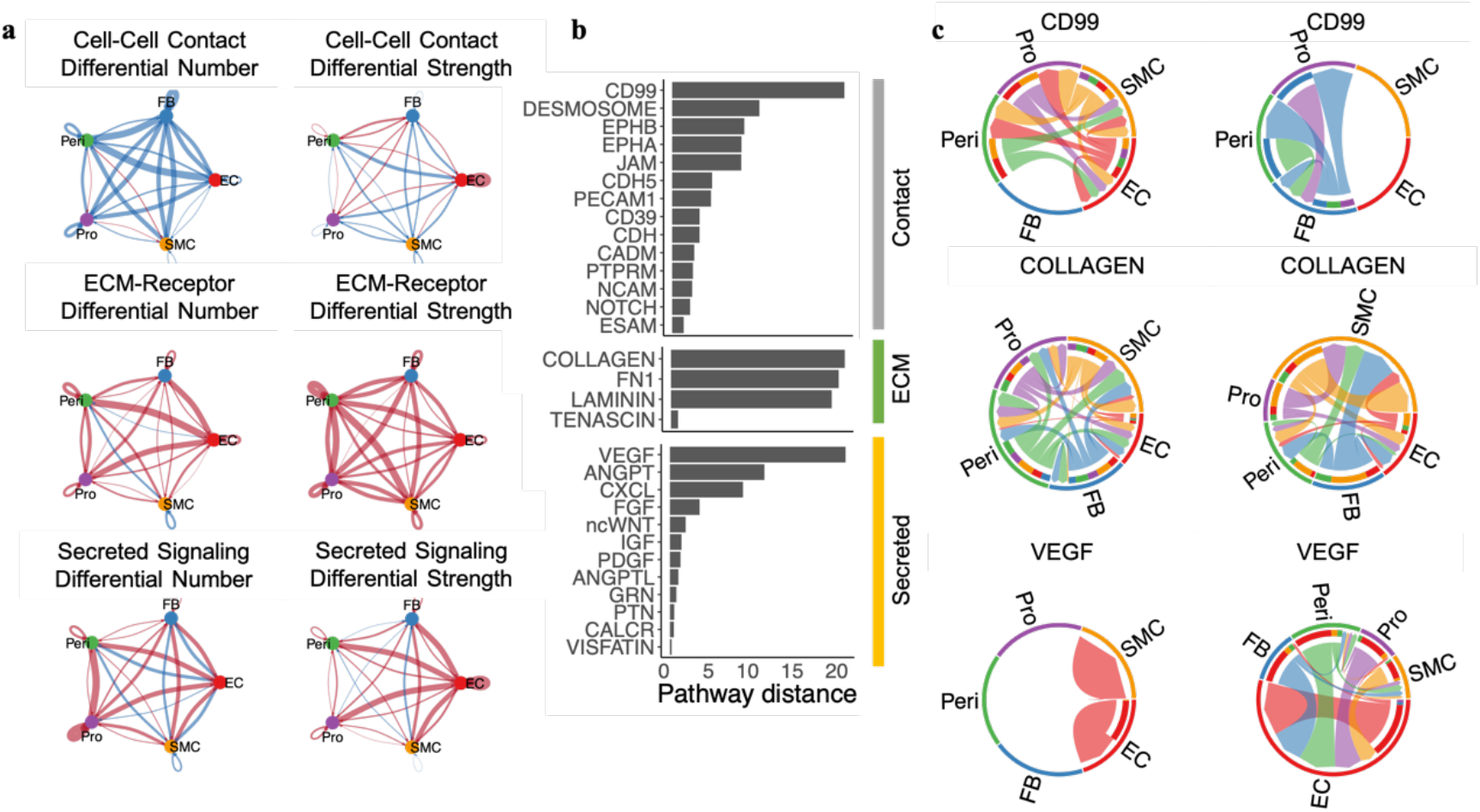
Variation in the intercellular communications. a) Differential number and strength of cell-cell contact, ECM receptor, secreted related signaling in 22q11.2 DS VO and control. The red (blue) edges in the circle plot represents the interaction number was enhanced (reduced). The width of red (blue) edges represented the enhanced (reduced) quantity. b) Distance of variable pathways associated with cell-cell contact, ECM receptor, secretion. c) Chord diagram of CD99, Collagen, CD99 signaling pathway network among cell types.

**Figure S7.**
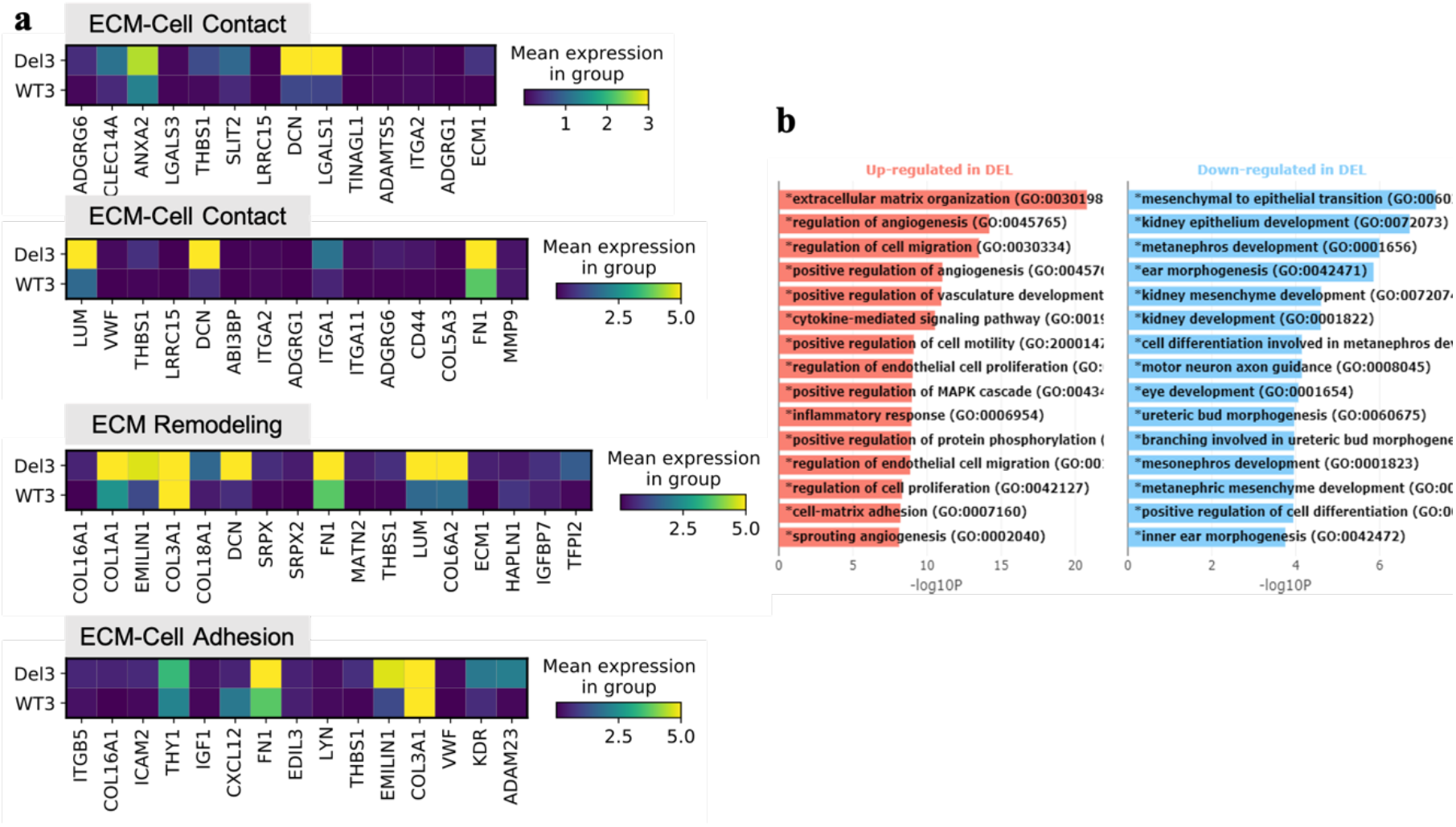
ECM-Cell interaction. a) Matrixplot of ECM-Cell contact, ECM remodeling, ECM-Cell adhesion associated gene expression in 22q11.2 DS VO and control. b) GO analysis of bulk RNA.

